# Pitfalls in re-analysis of observational omics studies: a post-mortem of the human pathology atlas

**DOI:** 10.1101/2020.03.16.994038

**Authors:** Jeroen Gilis, Steff Taelman, Lucas Davey, Lennart Martens, Lieven Clement

**Affiliations:** Department of Applied Mathematics, Computer Science & Statistics, Ghent University, Belgium; Data Mining and Modeling for Biomedicine, VIB Center for Inflammation Research, Ghent, Belgium; Department of Data Analysis and Mathematical Modelling, Ghent University, Belgium; VIB-UGent Center for Medical Biotechnology, VIB, Ghent, Belgium; Department of Biomolecular Medicine, Ghent University, Ghent, Belgium; Bioinformatics Institute Ghent, Ghent University, Ghent, Belgium

## Abstract

Uhlen *et al.* (Reports, 18 august 2017) published an open-access resource with cancer-specific marker genes that are prognostic for patient survival in seventeen different types of cancer. However, their data analysis workflow is prone to the accumulation of false positives. A more reliable workflow with flexible Cox proportional hazards models employed on the same data highlights three distinct problems with such large-scale, publicly available omics datasets from observational studies today: (i) re-analysis results can not necessarily be taken forward by others, highlighting a need to cross-check important analyses with high impact outcomes; (ii) current methods are not necessarily optimal for the re-analysis of such data, indicating an urgent need to develop more suitable methods; and (iii) the limited availability of potential confounders in public metadata renders it very difficult (if not impossible) to adequately prioritize clinically relevant genes, which should prompt an in-depth discussion on how such information could be made more readily available while respecting privacy and ethics concerns.

---

Uhlen *et al.* (*1*) published an open-access resource with cancer-specific prognostic marker genes based on a genome-wide transcriptome analysis of The Cancer Genome Atlas (TCGA) data compendium. For this study, they constructed more than 100 million Kaplan-Meier survival plots (Figure 1A), and corresponding significance levels were used to establish sets of favorable and unfavorable prognostic marker genes in different types of cancer. These markers were then communicated via the Human Pathology Atlas web resource (HPA). However, their data analysis workflow shows three shortcomings concerning specificity: (i) binarization of count data, (ii) multiple testing correction issues at two distinct levels, and (iii) ignoring baseline confounders. We here investigate the consequences and severity of these shortcomings, present more appropriate solutions for these problems, compare the results of our updated workflow to those of the original workflow and highlight challenges to prioritize clinically relevant genes from omics data of large-scale observational studies.

**Figure 1:**
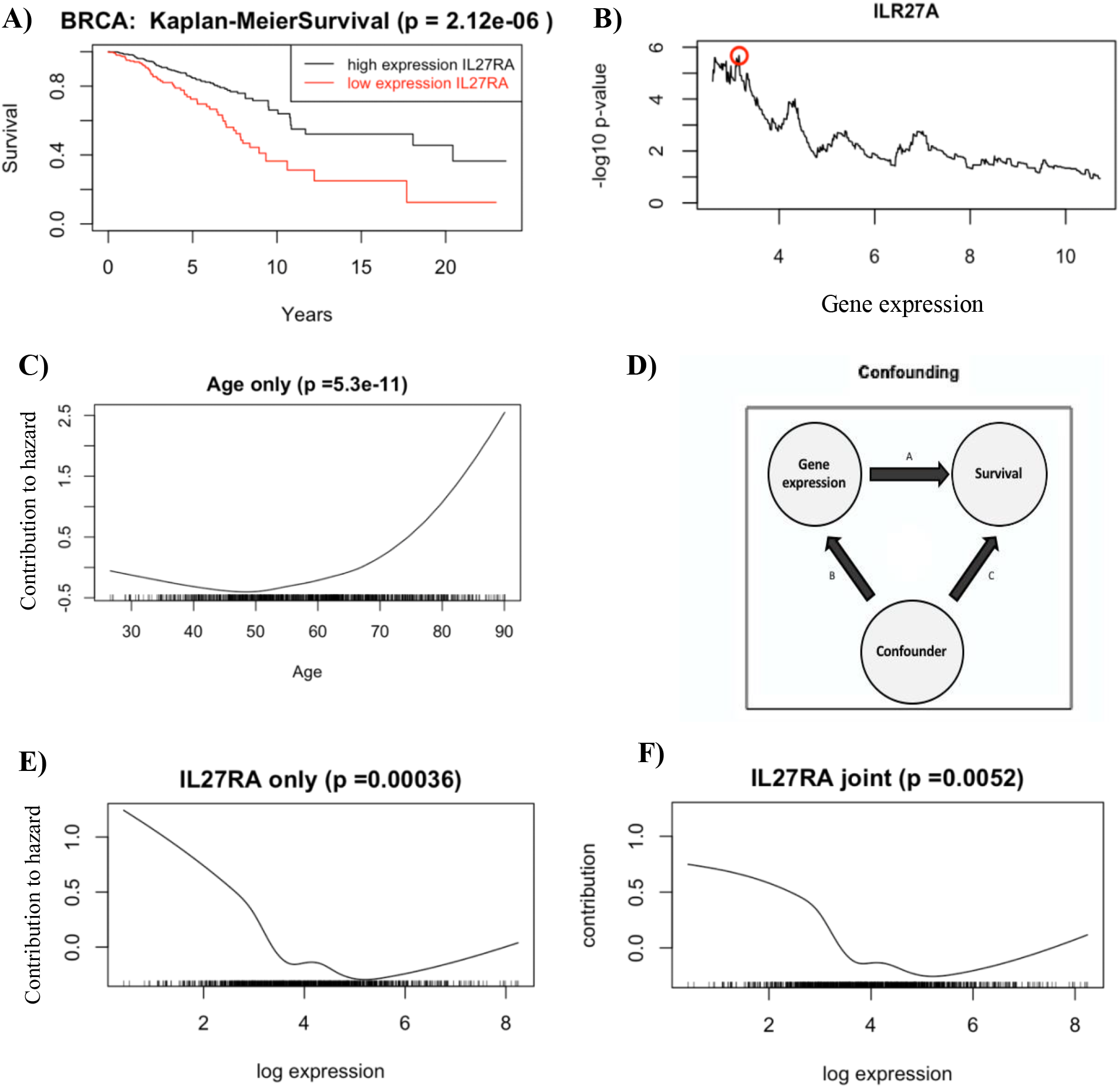
**A) Kaplan-Meier plot** for breast cancer patients stratified by the expression of the IL27RA gene (Uhlen et al., 2017 (1)). Patients with an IL27RA expression higher than the “optimal cut-off” display a higher survival rate. **B) Data dredging:** Distribution of log-rank p-values against the RNA expression of the IL27RA gene with different expression cut-offs on the BRCA dataset. A minimal p-value of 2.12E-06 is found at an expression threshold of 7.504. **C) A Cox model for age only** demonstrating the contribution of this confounder to the proportional hazard. Clearly, this hazard increases for increasing age. The flags at the bottom indicate observations. **D) Baseline confounders:** Confounders, such as a patient's age, affect both gene expression and survival. As a consequence, the effect of both arrows B and C could be misconstrued as a direct effect of gene expression. **E) A Cox model for the IL27RA gene** with its contribution to the proportional hazard as a function of its log expression. The flags at the bottom indicate observations. **F) A Cox model for IL27RA corrected for age effects**. As compared to figure 1E, the contribution of the IL27RA to the hazard has dropped for lower expression values and the p-value of the test is less significant (more than ten times higher). The flags at the bottom indicate observations.

## Binarization of count data

To create the 100 million Kaplan-Meier plots on which the study is based, patients are classified into two groups based on their gene expression (GE) level: above or below a certain cut-off value. This binarization leads to a huge loss of information, as information-rich GE count data are translated into the equivalent of an on/off state.

## Multiple testing issues

Of greater concern are the multiple testing issues, which occur at the level of the cut-off for binarization, and at the level of gene prioritization. Both are discussed below.

For the binarization, the criterion to bin GE levels was derived by exploring all possible GE cut-off values for each gene, thus generating hundreds of Kaplan-Meier plots *per* gene. Based on these plots, the cut-off with the best separation in survival between groups (i.e., the lowest p-value) was chosen (Figure 1B). The strategy thus *de facto* performs a large number of tests *per* gene, which requires correction for multiple testing. Yet such a correction was not performed and is moreover not trivial given the overall data analysis workflow and study design. As a result, there is a substantially increased risk for false positive marker selection. This can already be expected from the huge number of genes that were communicated as marker genes in e.g. the TCGA breast cancer dataset, with 847 genes having a p-value below 0.001 and 7749 genes with a p-value below 0.05, out of 17040 genes.

To demonstrate the repercussions of this data dredging strategy, we randomly shuffled the survival times from the TCGA database. This breaks the original association between the survival data and the GE profiles of the patients as well as the association between the survival data and baseline confounders. By applying the original binarization strategy adopted by Uhlen *et al.* (*1*) to this mock dataset, we obtained 320 genes with p-values below p<0.001 and 6410 genes with p<0.05, while a valid statistical procedure is expected to return around 17 (0.1% of 17040) and 852 genes (5% of 17040) under the null, respectively. Moreover, we show that the Uhlen *et al.* (*1*) method distorts the entire p-value distribution under the null hypothesis of no association (see Supplementary Materials Section 2.2.1).

When combining the results of all genes, a classical multiple testing problem arises. Indeed, thousands of genes are being tested as potential markers and multiple testing must be addressed. Again, no such correction is adopted in the analysis of Uhlen *et al.* (*1*).

Taken together, these two multiple testing problems, occurring at distinct two levels, create an intractable overall multiple testing problem, which inevitably leads to highly inflated false positive rates in the original Uhlen *et al.* analysis (*1*).

## Confounders

The third and final issue lies in the presence of confounders, which are not taken into account in the analysis of Uhlen *et al.* (*1*). For instance, in the TCGA breast cancer dataset, age has an extremely significant contribution to the hazard ratio (*P* = 5.3E-11, Figure 1C). As such, an analysis ignoring baseline confounders cannot correctly assess the impact of GE on survival (Figure 1 panels D - F).

An elegant solution to the three issues of the Uhlen analysis (*1*) is to build upon the semi-parametric Cox proportional hazards (CPH) framework (2) (see Supplementary Materials for details). Indeed, this allows us to model survival data using both categorical (e.g. gender) and quantitative predictors (e.g. GE, age, …), thus avoiding issues regarding the wasteful binarization of GE data and measured baseline confounders. For gene *IL27RA*, for instance, a decrease in hazard can be observed as GE levels increase (Figure 1F). Moreover, with the CPH approach we can assess the association between the expression of each gene and survival with a single test, in turn enabling the use of conventional false discovery rate (FDR) procedures to address the multiple testing problem (*3*).

## Analysis with the semi-parametric Cox proportional Hazards model

We illustrate the CPH framework on the TCGA breast cancer (BRCA) and liver cancer (LIHC) datasets. Note that all the results can be found on our companion GitHub page, https://github.com/statOmics/pitfallsOfHumanPathologyAtlas. We include a spline term for gene expression and for the available confounders (age in the BRCA analysis and age and body mass index in the LIHC analysis, see Supplementary Materials for details). For both datasets, we observe a clear discrepancy between the results provided by Uhlen *et al.* (*1*) and those of our improved workflow (Supplementary Tables 1–4).

For the BRCA dataset, including the confounder age strongly impacts the ranking of the genes, with many of the top genes originally identified moving to a substantially lower ranking in our analysis, and vice versa. Interestingly, several of the markers exclusively identified as high-ranking in our analysis are well-known breast cancer markers, for instance, *MMP13* and *RGL3*. This emphasizes the importance of correcting for confounders. Uhlen et al. (*1*) report 847 genes to be significantly associated with survival (uncorrected p-value < 0.001). Strikingly, no genes were found to be significantly associated with survival upon correcting for the confounder age and multiple testing (5% FDR significance level).

For the LIHC analysis, the impact on the ranking of the genes is comparatively small, as most genes in our top 10 list are among the 50 highest ranking genes in the original publication. As expected, the adjusted p-values obtained by our workflow are several orders of magnitude larger than those in the original publication, given that we appropriately correct for multiple testing. For the LIHC analysis, we identified 2812 genes that are significantly associated with survival (5% FDR significance level), markedly less than in the original publication (3501 genes with uncorrected p < 0.001).

Note, however, that the distribution of test statistics and p-values of all genes in our CPH approach indicates failure of the theoretical null distribution of the test statistic for both case studies (see Supplementary Materials). Efron (*4*) argued that this can be due to 4 reasons: failed mathematical assumptions, correlation across genes, correlation across patients, and unobserved confounders in observational studies. When we use his local false discovery rate approach, which exploits the massive parallel data structure to empirically estimate the null distribution, no significant association between gene expression and survival remains in both case studies. We also further examined why the theoretical null distribution failed in our context, and could show that the semi-parametric Cox model suffered from all of the above four reasons in both case studies (see Supplementary Materials). However, when replacing the spline-based likelihood ratio test (LRT) with an LRT for a linear and quadratic GE-effect, we no longer observed failure of the null due to mathematical assumptions. Nevertheless, the null of this quadratic CPH model still suffers from unobserved confounders (BRCA and LIHC) and from correlation between genes (LIHC). The former, however, could not be addressed using the publicly available baseline confounders. This indicates the challenges of prioritizing clinically relevant genes from large scale observational studies.

## Conclusion

The advent of large open-access, knowledge-based efforts, such as TCGA, opens the way for high-throughput statistical (re-)analyses to facilitate novel experiments, e.g. through the construction of prioritized gene sets. However, our assessment highlights that there are at least three distinct problems with such datasets today: (i) re-analysis results can not necessarily be taken forward by others; (ii) current methods are not necessarily optimal for the re-analysis of such data; and (iii) the limited availability of potential confounders in the public metadata renders it very difficult (if not impossible) to adequately prioritize clinically relevant genes. Possible solutions to these problems are to (i) involve statisticians in such large-scale re-analyses; (ii) to foster further developments in causal inference, survival analysis and multiple testing for large-scale omics analyses from observational studies; and (iii) to develop ways to make potential confounders available for such large-scale, open-access data compendia while adhering to relevant privacy and confidentiality concerns.

## Acknowledgements

LC and JG are supported by the Research Foundation Flanders (FWO), research grant No. G062219N, and JG is further supported by FWO SB fellowhip No. 3S037119. LM is supported by the FWO research grant No. G042518.

## Supplementary material

This document serves as supplementary material for the technical comment on “A pathology atlas of the human cancer transcriptome” by Uhlen *et al.*, 2017 (*1*), describing the rationale and methodology of each step in our revised workflow, supplemented with R-code on our GitHub companion page: https://github.com/statOmics/pitfallsOfHumanPathologyAtlas. Analyses will be limited to the breast cancer (BRCA) and liver cancer (LIHC) datasets supplied by The Cancer Genome Atlas (TCGA) project via the Genomic Data Commons (GDC) data portal, but the procedures for other datasets in the TCGA repository are analogous.

## 1. Reconstruction of the original Kaplan-Meier analysis

First, we reconstruct the Kaplan-Meier plots from the survival analysis of the original publication by Uhlen *et al.* (*1*). There, patients were divided in two groups based on the expression value of a certain gene, i.e. above or below a certain cut-off value. To gain insight in the rationale of their methodology, we reconstruct these plots for the BRCA dataset (Figure S1). For concordance with the original publication, our analysis uses FPKM (fragments per kilobase million) expression values. Here, we only show the Kaplan-Meier curves of the top two favorable and unfavorable prognostic marker genes.

**Figure S1:**
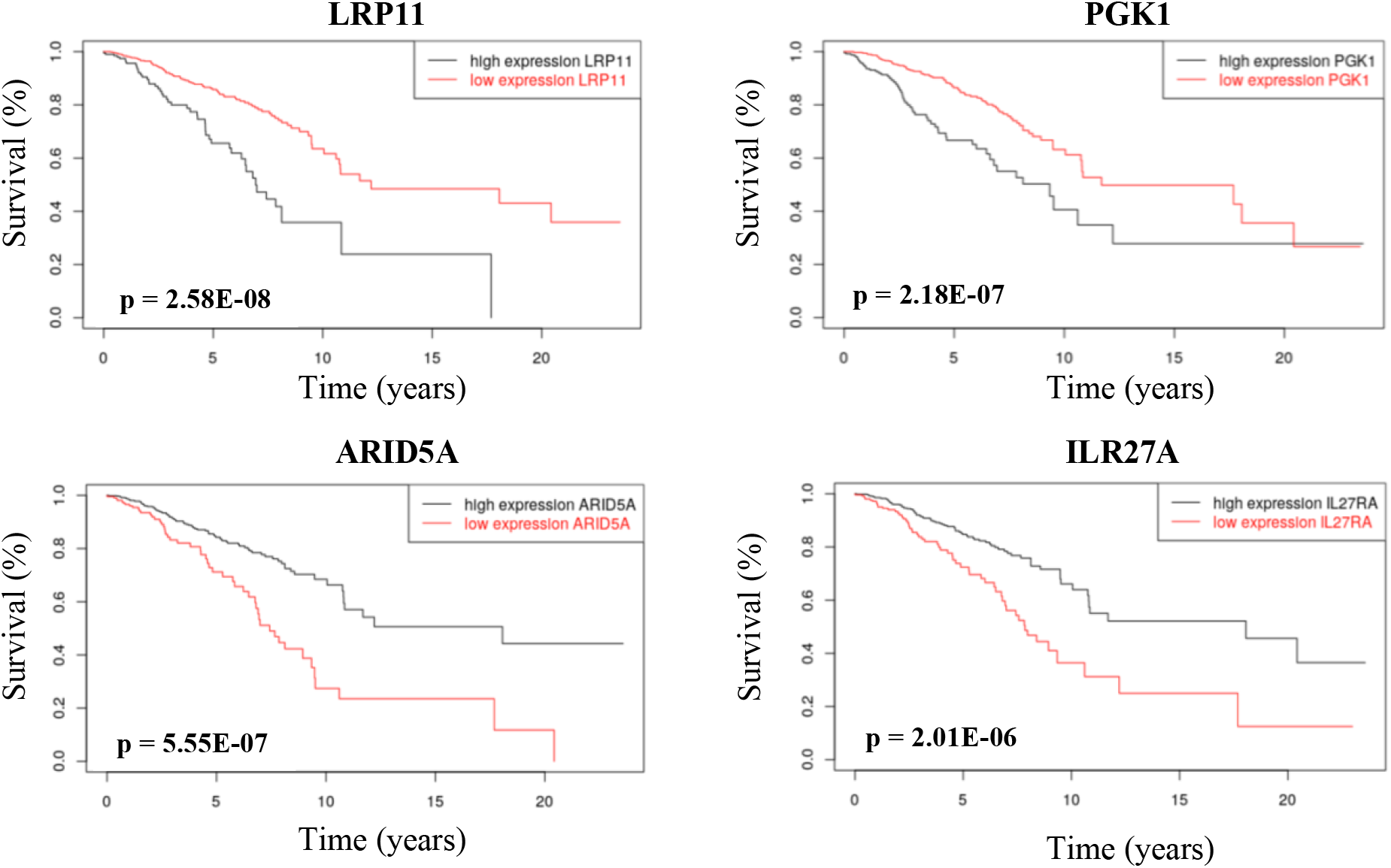
Example of Kaplan-Meier plots for two of the most significant favorable markers (top row) and unfavorable markers (bottom row) in the TCGA breast cancer dataset. The expression of the selected genes has a strongly significant association with survival based on the Kaplan-Meier analysis. The p-values are indicated in the plot. Note that these values might slightly deviate from those in the original publication, due to updates in the TCGA data compendium. Black and red lines show the survival of patients with gene expression values higher and lower than the optimal cut-off value, respectively.

The source code for creating these plots is available on our companion GitHub page /BRCA/originalBRCA.Rmd. Note, that the resulting p-values and plots might not be identical to those published by Uhlen *et al.* (2017), due to regular updates of the TCGA data compendium.

## 2. Shortcomings of the original Kaplan-Meier analysis

We report three shortcomings of the original Kaplan-Meier analysis concerning specificity of the reported prognostic marker genes: (i) binarization of count data, (ii) multiple testing correction issues at two distinct levels, and (iii) ignoring baseline confounders. Here, we will discuss each of these shortcomings in depth. We first show where the existing issues cause problems in the analysis, which we subsequently prove using simulated data. Next, we proceed to correct for these issues using appropriate methods in a revised and more reliable workflow.

### 2.1. Binarization of count data

To regenerate the original Kaplan-Meier plots, patients are classified into two groups based on their gene expression level: above or below a certain cut-off value. This binarization leads to a huge loss of information, as information-rich gene expression count data are translated into the equivalent of an on/off state. This wasteful binarization of the expression values, however, can be avoided altogether by modeling survival using semi-parametric Cox proportional hazards models (2), which we will discuss in Section 3.

### 2.2. Multiple testing problems and p-hacking

The cut-offs on the gene expression values to generate Figure S1 were obtained from the supplementary tables of the original publication. In order to achieve the largest possible distinction between the survival of both patient groups, these cut-offs were generated with a data dredging strategy. More explicitly, the dataset was exhaustively searched for gene-expression cut-offs (i.e. prior to the actual data analysis) which would finally lead to the smallest possible p-value for each gene. The source code for generating these plots for the breast cancer dataset is available on our companion GitHub page /BRCA/originalBRCA.Rmd. The plot for the *IL27RA* gene in the BRCA dataset is shown in Figure S2.

**Figure S2:**
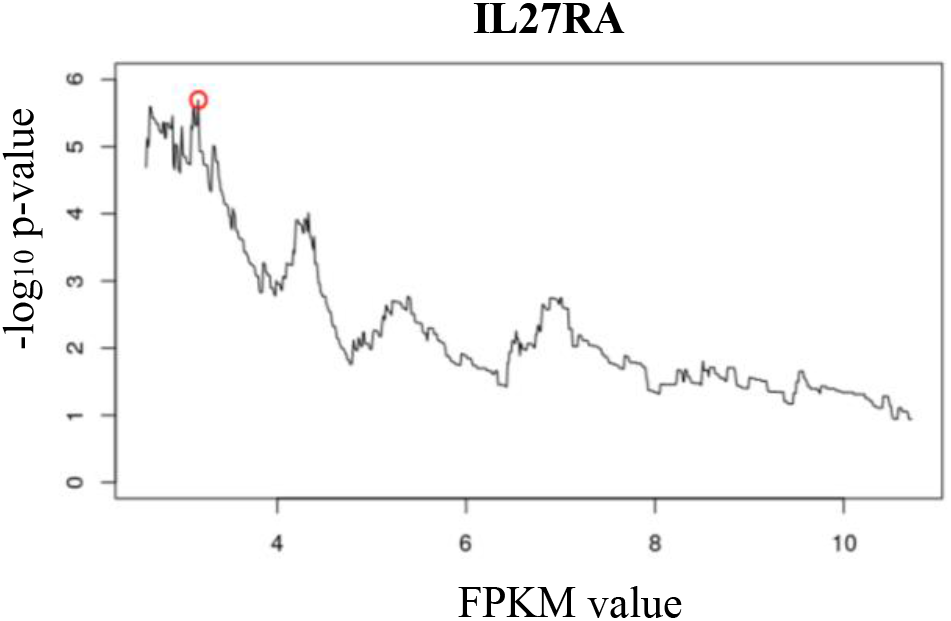
Selecting a cut-off gene expression value for the gene *IL27RA* as described by Uhlen *et al.* The cut-off value is selected by evaluating all gene expression (FPKM) values from the 20_th_ to the 80_th_ percentile and by subsequently selecting the value that achieves the most significant difference in the survival of the resulting groups. Significance is expressed as the lowest Kaplan-Meier p-value (highest −log10 p-value). Note that the calculated cut-off values might not be identical to those supplied by Uhlen *et al.* in 2017 due to recent updates of the TCGA data compendium.

This strategy thus *de facto* performs a large number of tests *per* gene, which requires correction for multiple testing. Yet, such a correction was not performed and is moreover not trivial given the overall data analysis workflow and study design. As a result, there is a substantially increased risk for false positive marker selection. This can already be expected from the huge number of genes that were communicated as marker genes in e.g. the TCGA breast cancer dataset, with 847 genes having a p-value below 0.001 and 7749 genes with a p-value below 0.05, out of 17040 candidate genes. For the LIHC dataset, the original publication even suggested as much as 3501 genes with a p-value below 0.001 and 10632 genes with a p-value below 0.05, out of 16397 candidate genes.

Below, we will make use of simulated data to demonstrate the repercussions of this data dredging strategy.

#### 2.2.1. Mock analysis

To demonstrate the repercussions of the data dredging strategy, we randomly shuffled the TCGA survival data (companion GitHub page /BRCA/originalBRCA_problems.Rmd). This breaks the original association between the survival data and the GE profiles as well as the association between the survival data and baseline confounders. Hence, this mock dataset does not contain marker genes that are associated with survival. By applying the original binarization strategy adopted by Uhlen *et al.* (*1*) to this mock dataset, however, we obtained 320 genes with p-values below p<0.001 and 6410 genes with p<0.05, while a valid statistical procedure is expected to return 17 (0.1% of 17040) and 852 genes (5% of 17040) under the null, respectively. In Figure S3, we show Kaplan-Meier plots that were generated with Uhlen *et al.* strategy for the top genes of the mock analysis.

**Figure S3:**
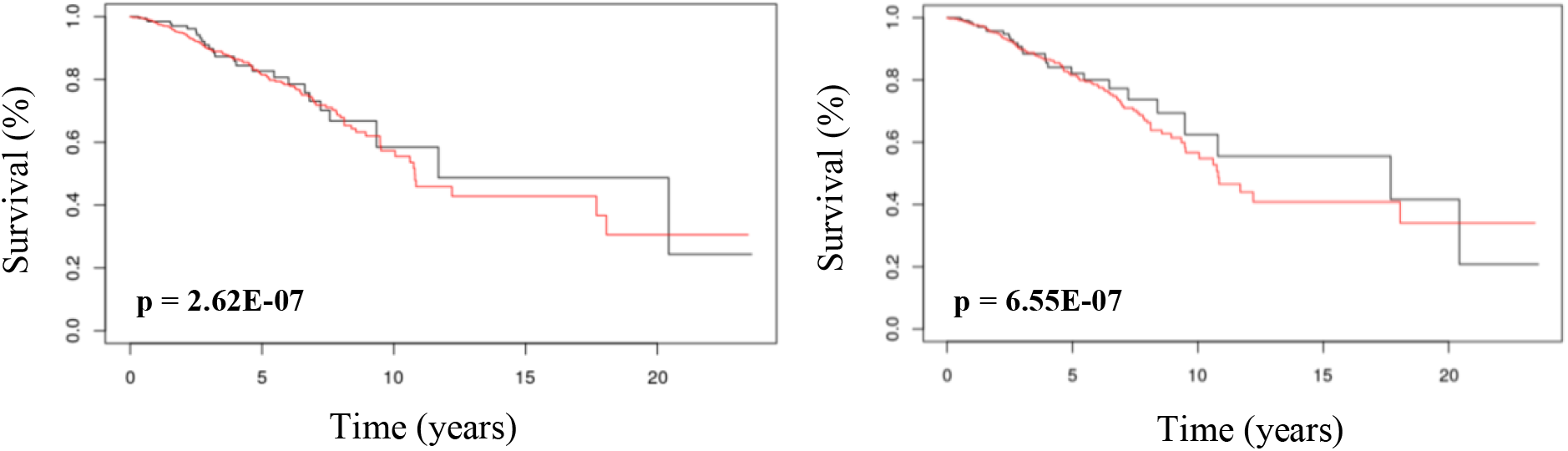
Example of Kaplan-Meier plots for two of the most significant genes in the mock analysis. With the cut-off-based strategy that was adopted in the original publication of Uhlen *et al.* (2017), p-values as small as 2.62E-07 were obtained in the mock dataset for which no significant association between gene expression and survival occurs.

Figure S4 (left panel) shows that the distribution of the p-values is unnaturally skewed towards small values; strikingly, the largest p-value that was obtained for this mock dataset, which again does not contain markers, was p = 0.65. In principle, valid p-values under the null hypothesis should be uniformly distributed over the entire [0,1] interval. As such, it becomes clear that the results reported by the Uhlen *et al.* method are meaningless.

Intriguingly, the distribution of the p-values in the original Kaplan-Meier analysis for the BRCA dataset are equally skewed towards small values (Figure S4, right panel). For a valid statistical procedure, one would expect a uniform distribution of p-values in the absence of prognostic marker genes. The presence of a number of genes that are truly associated with survival would result in an inflation of p-values close to zero, however, the larger p-values are supposed to be uniformly distributed. The observed pattern clearly demonstrates that the resulting p-values of the Uhlen *et al.* analysis are strongly biased towards small values as a consequence of the data dredging strategy for determining the binary gene expression cut-off values.

**Figure S4:**
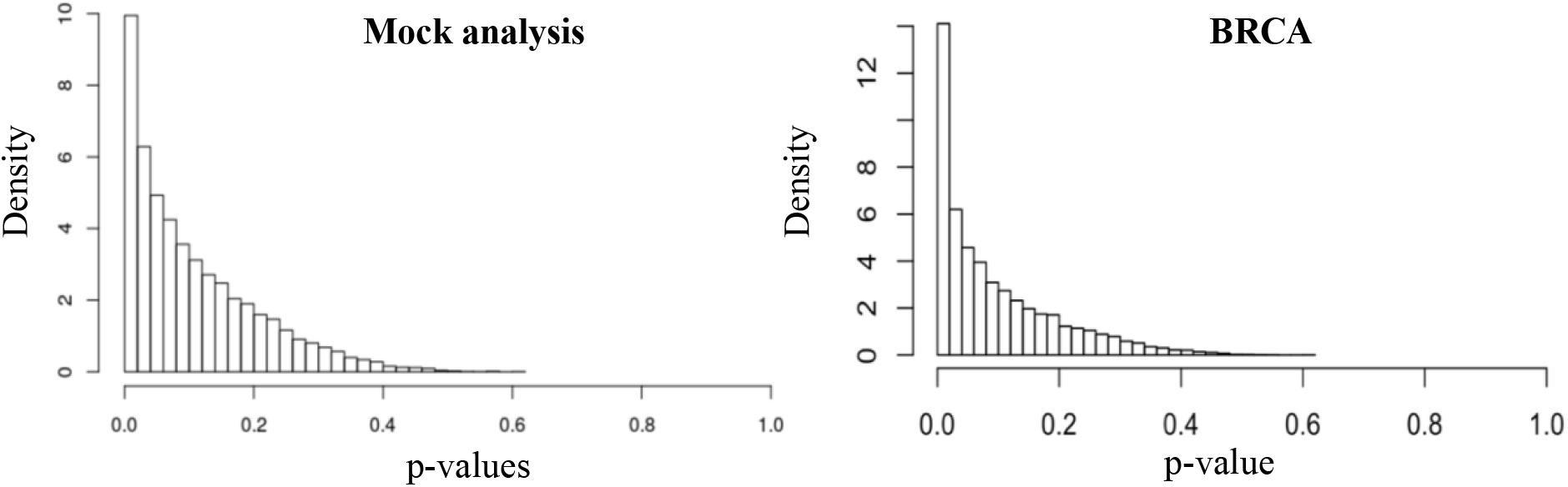
Histogram of the distribution of the Kaplan-Meier p-values in the mock analysis. **Left panel**; As a consequence of the p-hacking strategy, no values larger than p = 0.65 are obtained, even in the analysis of the mock dataset. While under the null hypothesis the p-values of a valid statistical inference procedure are expected to be uniformly distributed over the [0,1] interval (i.e. no association between any of the genes and survival), their observed distribution is strongly skewed towards small values. **Right panel**; When displaying the p-values of the original publication for the BRCA dataset, we can see that they are equally skewed towards small values. This clearly demonstrates that the resulting p-values of the original analysis are strongly biased towards small values as a consequence of the p-hacking strategy, which implies an accumulation of false positive markers.

### 2.3. Confounders

As a third shortcoming, the original analysis ignores baseline confounders. Confounders disrupt the causal interpretation of the dependent and explanatory variable(s). Since factors such as the age of a patient already have significant impact on survival and are also influencing the expression of certain genes, the analysis should be corrected for such confounders when assessing the impact of gene expression on survival. The earlier proposed Cox proportional hazards model also allows to include confounders and shall hence be used for a more reliable survival analysis, as discussed in Section 3.

## 3. Cox proportional hazards models

We propose the use of the semi-parametric Cox proportional hazards (CPH) models for revisiting the original analysis of the TCGA datasets (*2*). First, CPH models circumvent binarizing gene expression values, as they allow for the modeling of quantitative variables, directly. Second, treating the gene expression value as a quantitative variable removes the need for exhaustively searching the dataset for artificial gene expression cut-off values. This greatly eases the downstream multiple testing correction. Indeed, we now only need to correct for the fact that multiple genes were assessed simultaneously, for instance, by adopting the false discovery rate procedure as described by Benjamini and Hochberg (*3*). Finally, CPH models allow for confounders to be included as covariates. Hence, the analysis will be more reliable with respect to finding marker genes that are truly associated with the cancer phenotype.

The Cox hazard function can be interpreted as the risk of undergoing a certain event (in this case, death) at time t:

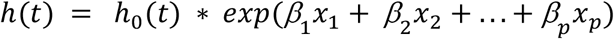

In this equation,

- h_0_(t) is the baseline hazard (which can vary throughout time)
- X is a set of covariates each with an effect size β. In this CPH model X captures the association of the expression for the gene in question. To correct for confounders, additional covariates such as age or body mass index (BMI) can be incorporated.

In order to achieve a good fit and to capture potential nonlinear associations, we use a spline function. Spline functions expand a quantitative predictor (z) in a set of basis functions: e.g. x_1_ … x_k_:

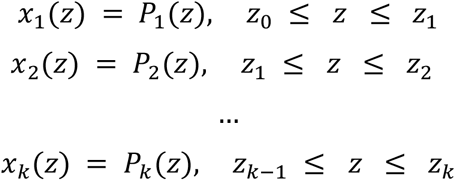

In this equation,

- *P*_*i*_(*z*) are different polynomials for gene expression value *z*
- The given k+1 points are referred to as knots, these can be evenly spaced throughout the parameter space like in the *pspline*-function of the Survival R package (*4*). Due to the nature of the observational study, however, the predictor data are much denser in intervals in the middle than intervals at the boundaries. To avoid overfitting in the boundary regions the original *pspline*-function is modified to a user-defined spline function referred to as ‘*qspline’*, where knots are set at evenly spaced quantiles. This made the fit more robust against outliers and modelled the middle of the predictor space with increased detail.

Note; for all analyses that are not reconstructions of the work by *Uhlen et al.* (*1*), we work with log-transformed counts-per-million as gene expression values. As the expression data is dissected for each gene and not across all genes, FPKM expression values do not provide added value to the analysis. All analyses related to the CPH models can be retrieved on our compagnion GitHub repository, i.e. from scripts /BRCA/revisitedBRCA.Rmd and /LIHC/revisitedLIHC.Rmd for the breast cancer and liver cancer datasets, respectively.

## 4. Deviations from the theoretical null distribution

We applied CPH models to quantify the strength of the association between the expression of each gene in the dataset and patient survival, after correcting for patient specific confounders. In brief, a likelihood ratio test between two models was used, comparing the fit between (i) a CPH model that models a patient’s survival based only on this patient’s age (BRCA dataset) or age and BMI (LIHC dataset), and (ii) a CPH model that models survival based on the confounders and gene expression values. Note, that we use splines to model the association with gene expression and the confounder(s).

In transcriptomics data analysis, it is commonly assumed that most genes in the dataset are null genes, i.e. most genes are not associated with survival. Under the assumption of only null genes, the distribution of the p-values, that quantify the significance of the association between gene expression and survival, are uniformly distributed over the [0,1] interval. The presence of a relatively small number of genes that are truly associated with survival will result in an inflation of p-values close to zero. As shown in the right panel of Figure S4, the data dredging strategy that was adopted in the original publication leads to a p-value distribution that is strongly skewed towards small values. However, when taking a closer look to the p-values derived from our proposed workflow for BRCA dataset, an almost linear decrease over the entire [0,1] interval is observed (Figure S5, left panel). While for the LIHC dataset there seems to be a clear inflation of near-zero p-values, a linear decrease in the distribution of p-values in the [0.1, 1] interval can still be observed (Figure S5, right panel).

**Figure S5:**
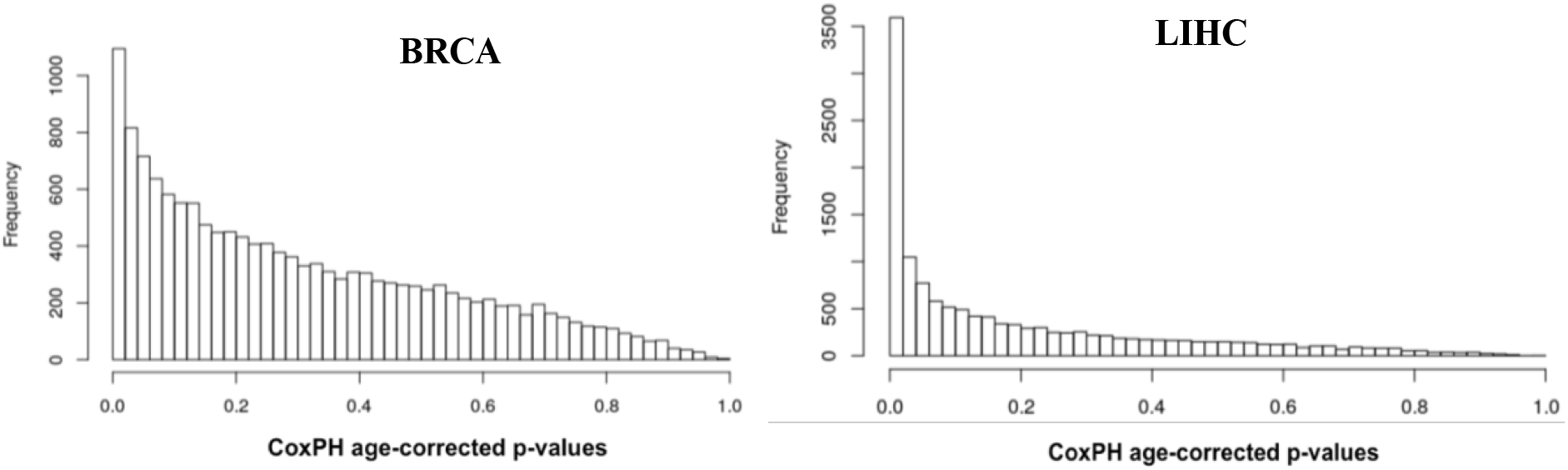
Histogram of the distribution of the p-values for our proposed Cox proportional hazards model. **Left panel**; p-values for the breast cancer datasets. Under the null hypothesis, i.e. when assuming no association between any of the genes and survival, a uniform distribution of p-values is expected. When association between some of the genes and survival is present, this would result in an inflation of near-zero p-values. Here, in contrast, the distribution of the p-values seems to decrease almost linearly over the [0,1] interval, which suggests a violation of an underlying assumption of the null distribution of the test statistics. **Right panel**; p-values for the liver cancer dataset. While for this dataset there seems to be a clear inflation of near-zero p-values, a linear decrease in the distribution of p-values in the [0.1, 1] interval can still be observed.

Such behavior indicates failure of the theoretical null distribution. In this example, one would expect the likelihood ratio test statistic to be chi-squared distributed under the null hypothesis, with the number of degrees of freedom equal to those of the CPH models (which were set to df = 4 in our analysis, see scripts /BRCA/revisitedBRCA.Rmd and /LIHC/revisitedLIHC.Rmd) and we expect the majority of the genes to follow the null. However, this assumption is clearly not met for the majority of the test statistics in our respective workflows (see Figure S6).

**Figure S6:**
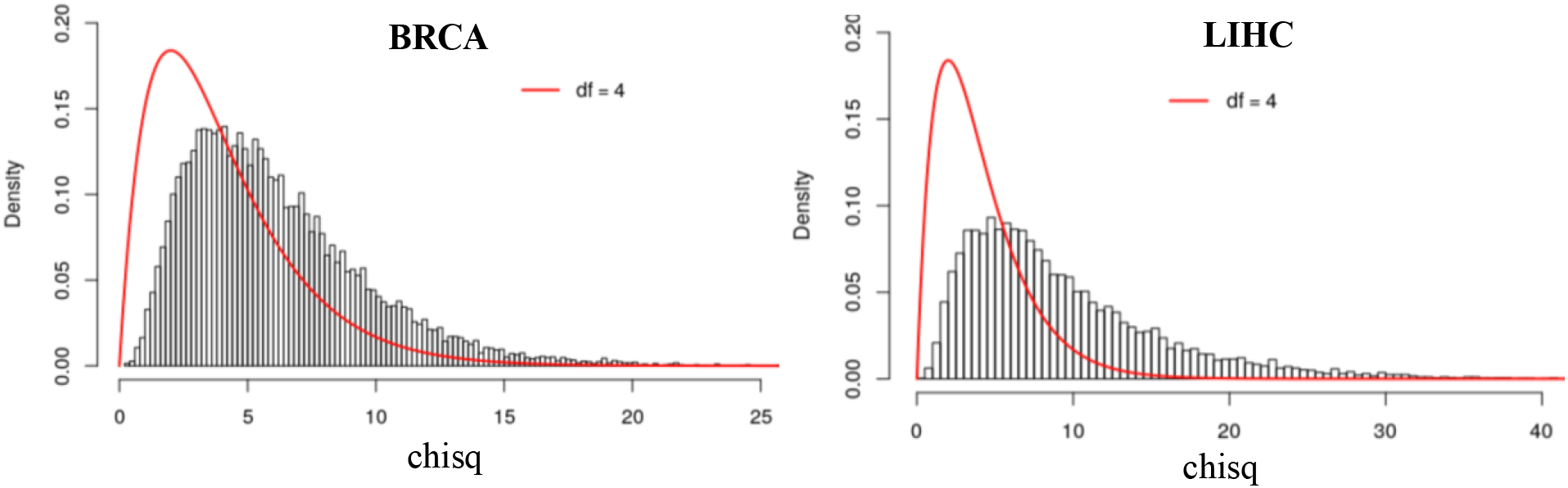
The majority of the test statistics do not follow the theoretical null distribution. **Left panel**; breast cancer dataset. Under the null hypothesis, the test statistics are expected to follow a chi-squared distribution with 4 degrees of freedom (red line). However, the test statistics are shifted towards larger values, indicating that a massive number of genes are associated with survival or failure of the theoretical null jeopardizing reliable inference. **Right panel**; liver cancer dataset. A similar behavior in the distribution of test statistics as in the breast cancer dataset is observed.

Efron (*5*, *Chapter 6*) describes four reasons why the theoretical null distribution may fail; (I) failed mathematical assumptions, (II) correlation across genes, (III) correlation across patients, and (IV) unobserved confounders in observational studies. A schematic representation of the data structure in our study is depicted in Supplementary Figure S7 and illustrates reasons (II), (III) and (IV).

**Figure S7:**
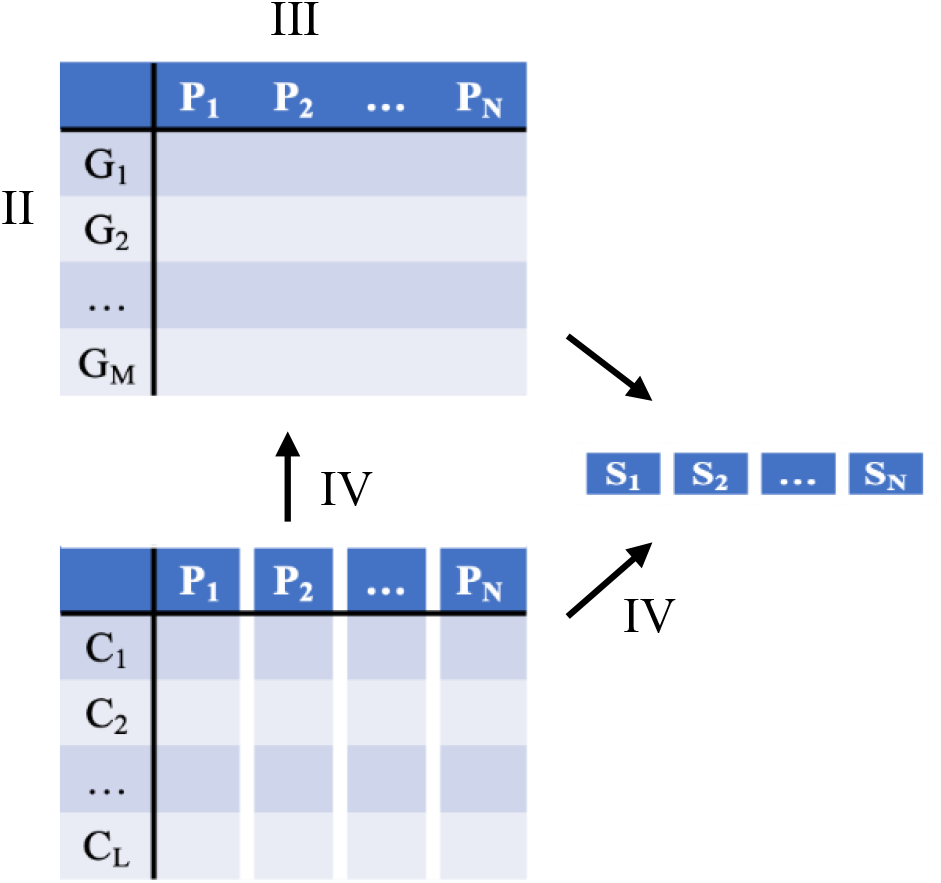
Schematic representation of the data structure in this study. The response variable, survival time (S), is a vector of length *N*, with *N* the number of patients in the BRCA or LIHC datasets. There are two matrices of explanatory variables. On the top left, we have an *M* × *N* matrix of gene expression values, with *M* the number of genes. On the bottom left, we have an *L* × *N* matrix of confounders, with *L* the number of confounders. Note, that these confounders may be measured or unmeasured. The arrows indicate dependencies between the different data types. We also indicate three of the reasons discussed by Efron (*5*) for failure of the theoretical null; (II) correlation across genes, (III) correlation across patients and (IV) unmeasured confounders. Cells and columns that are separated suggest independent observations, whereas concatenated cells suggest dependent or correlated observations.

Efron argues to correct for these issues by estimating the null distribution empirically. Particularly, we will make use of the companion ‘locfdr’ package developed by Efron et al. (*6*). As suggested by Efron (*5*, *Chapter 6*) for test statistics that follow a chi-squared theoretical null distribution, we first convert the p-values obtained from the analysis of the BRCA and LIHC datasets, respectively, to z-scores according to,

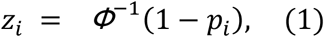

with p_i_ the original p-value indicating the significance of association between the expression of gene *i* and survival, Φ the cumulative distribution function for the standard normal distribution and zi the resulting z-score for gene *i*. Note, that for this transformation associations between gene expression and survival with small p-values all will end up in the right tail of the z-score distribution. We now visualize the resulting z-score distributions for the BRCA and LIHC datasets, respectively (Figure S8, panels A and B).

Efron (*5*), further, assumes a mixture distribution for the z-scores, i.e.

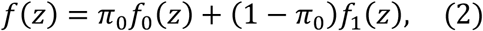

with *f*(*z*) the mixture distribution of the z-scores, *f*_0_(*z*) the distribution of the z-scores for the genes that are not associated with survival (under *H*_0_) and *f*_1_(*z*) the distribution of the z-scores for the genes that are associated with survival (under *H*_1_) and *π*_0_ the proportion of genes that are not associated with survival. Efron then defines the local false discovery rate (lfdr) as

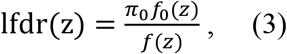

i.e. the posterior probability that a specific gene with a score z is a null gene (false positive) (*7*). He also has shown the link between the lfdr and the conventional FDR, which is the expected number of false positives (null genes) in the set of significant genes, say *S*, that is returned. In fact, the FDR is the expected value of the lfdr of all genes in set *S*,

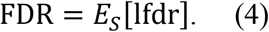

He also argues that the use of an lfdr significance level of 0.2 approximately corresponds to an FDR significance level of 0.05 in many applications. With his package locfdr (*6*), *π*_0_, *f*_0_(*z*) and *f*(*z*) are estimated empirically by exploiting the massive parallel structure of omics data.

For the BRCA dataset, the empirical null distribution of the resulting z-scores is shifted as compared to the theoretical standard normal distribution (Δ = 0.656). For the LIHC dataset, the resulting z-scores are shifted as well (Δ = 1.163), and the distribution additionally becomes wider than the theoretical null (σ = 1.217). As such, it is clear that inference based on the theoretical null distribution would lead to erroneous results. If we instead make use of the lfdr approach proposed by Efron et al. (*6*, *7*), no significant associations between gene expression and survival remain in either case study (lfdr < 0.2).

However, we do observe that the estimated empirical null component densities (blue dashed curves in Figure S8, panels A and B) do not seem to fit the observed distribution of z-scores very well. In both the BRCA and LIHC datasets, there are clear deviations between the estimated null component density and the observed z-score distribution in the left tail of the distribution. The deviations in the right tail of the distribution for the LIHC dataset (Figure S8 panel B) are harder to interpret, as they may either arise from a suboptimal fit of the empirical null or the presence of genes that are truly associated with survival. Note that errors in fitting the null density in the right tail are more severe than errors in the left tail from a practical perspective, as genes that are associated with survival will all be present in the right tail, given how the z-scores were generated (Formula 1).

To further examine why the theoretical null distribution fails for these two datasets, we first omit the correction for the baseline confounders (Figure S8, panels C and D). This did not have a noteworthy impact on the estimation of the empirical null distribution, suggesting that correcting for confounders does not have a strong impact on the estimation of the empirical null. Next, we model the association between gene expression and survival with a linear and a quadratic term for gene expression (Figure S8, panels E and F), rather than spline term (Figure S8 panels A-D). For the BRCA dataset, the empirical null distribution of the resulting z-scores remains shifted as compared to the theoretical standard normal distribution (Δ = 0.412) and has additionally become wider (σ = 1.151). For the LIHC dataset, the resulting z-scores are also shifted (Δ = 0.930) and wider than the theoretical null (σ = 1.456). The imperfect fit of the empirical null to the left tail of the observed z-score distribution remains for the LIHC dataset but seems to be resolved for the BRCA dataset.

**Figure S8:**
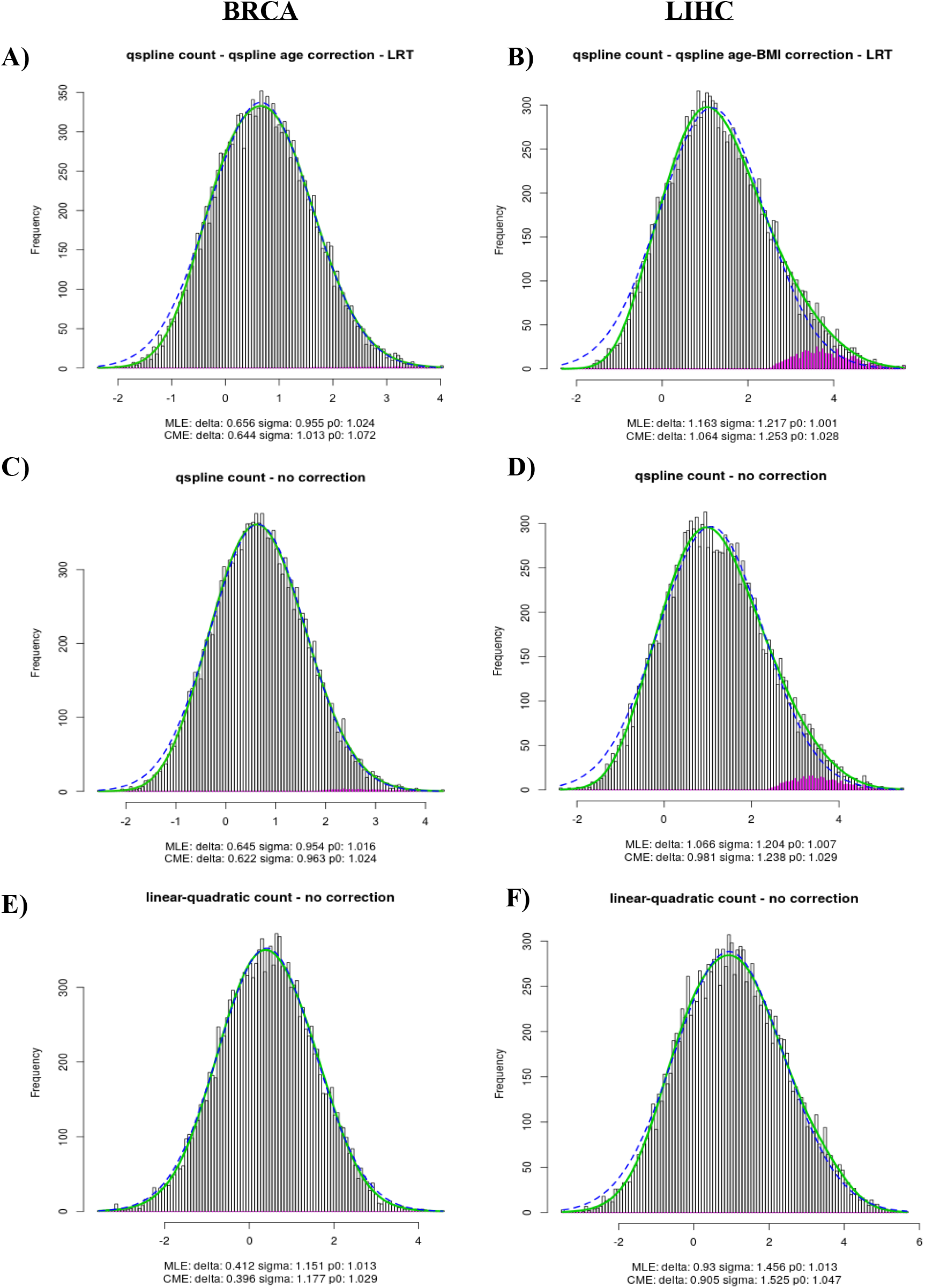
Z-scores for the association between gene expression and survival on the original BRCA (left panels) an LIHC (right panels) datasets. The blue dashed curves represent the estimated empirical null component densities. The green solid curves represent the fitted mixture density, i.e. a mixture between the null genes and marker genes (non-null). **Panels A and B; Z-score distributions when modelling gene expression with a spline function, while correcting for confounders**. For the BRCA dataset, the empirical null distribution of the resulting z-scores is shifted as compared to the theoretical standard normal distribution (Δ = 0.656). For the LIHC dataset, the resulting z-scores are shifted as well (Δ = 1.163), and the distribution additionally becomes wider than the theoretical null (σ = 1.217). Note that the empirical null distribution does not very well fit the observed z-scores in the left tail of the distribution. The deviations in the right tail of the distribution for the LIHC dataset (Figure S8 panel B) are harder to interpret, as they may either arise from a suboptimal fit of the empirical null or the presence of genes that are truly associated with survival. **Panels C and D; Z-score distributions when modelling gene expression with a spline function, ignoring confounders**. As compared to panels A and B, we here did not correct the analysis for confounders. This only had a very minor impact on the estimation of the empirical null distribution. **Panels E and F; Z-score distributions when modelling gene expression with a linear and a quadratic term, ignoring confounders**. For the BRCA dataset, the empirical null distribution of the resulting z-scores remains shifted as compared to the theoretical standard normal distribution (Δ = 0.412) and has additionally become wider (σ = 1.151). For the LIHC dataset, the resulting z-scores are also shifted (Δ = 0.930) and wider than the theoretical null (σ = 1.456). The imperfect fit of the empirical null to the left tail of the observed z-score distribution remains for the LIHC dataset but seems to be resolved for the BRCA dataset.

Next, we systematically assess which of four potential reasons is involved in the failure of the theoretical null distribution in both our case studies.

The issue of failed mathematical assumptions in our applications would mean that the asymptotic chi-squared distribution is invalid for instance due to the violation of the proportional hazards assumption or because of the use of penalized regression with splines.

As described by Efron (*5*), the issue of failed mathematical assumptions is the only type of failure that can be addressed via permutations. In the remainder of this section we will exploit permutation strategies to generate data under the null hypothesis of no association between gene expression and survival under controlled assumptions. So, the analyses displayed below can be solely used to identify potential reasons for failure of the null distribution and not to improve the analysis as such.

Under the first strategy we randomly permute the gene expression data between patients for each gene separately (Figure S9). This breaks the correlation between different genes within patient as well as the association with both survival and the confounders. Hence, under this permutation strategy, the empirical null distribution can only deviate from the theoretical null distribution if there is a violation of the assumptions of the underlying mathematical model.

**Figure S9;.**
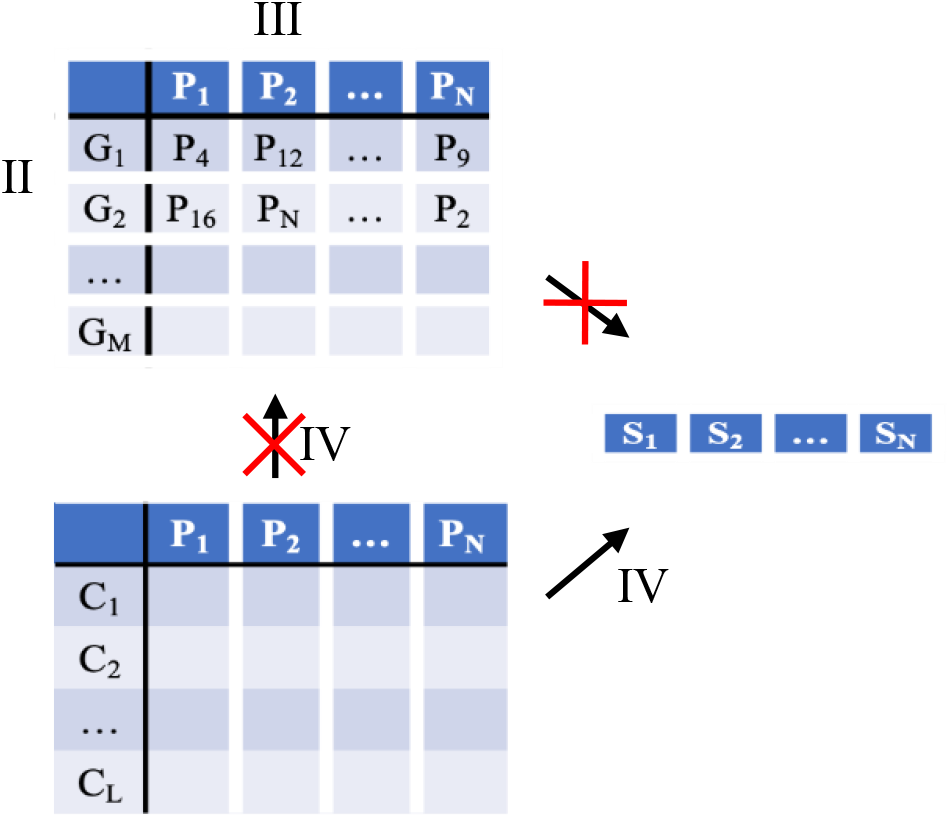
First permutation strategy. We randomly permute the gene expression data between patients, for each gene separately. This breaks the correlation between different genes within patient (issue II), as well as the association between gene expression on the one hand and survival and the confounders on the other hand, as indicated by the red crosses. Hence, under this permutation strategy, the empirical null distribution can only deviate from the theoretical null distribution if there is a violation of the assumptions of the underlying mathematical model (issue I).

The empirical null distribution for one permuted dataset where the expression of each gene is shuffled independently is depicted in Figure S10. When assessing the association between gene expression and survival with a spline-based likelihood ratio test, shifts and deviations in the width of the empirical null distribution as compared to the theoretical null remain in the both the permuted BRCA and LIHC datasets, suggesting failed mathematical assumptions. By contrast, when the analysis is performed with a likelihood ratio test based on a model with a linear and quadratic gene expression effect, we no longer observe failure of the null due to mathematical assumptions; the estimated empirical null distribution overlaps almost perfectly with the theoretical standard normal distribution upon conversion of the p-values in z-scores.

The failure of the mathematical assumptions of the spline-based model may be explained by the fact that we adopted penalized estimation and further research is required to efficiently unlock splines within our context.

**Figure S10:**
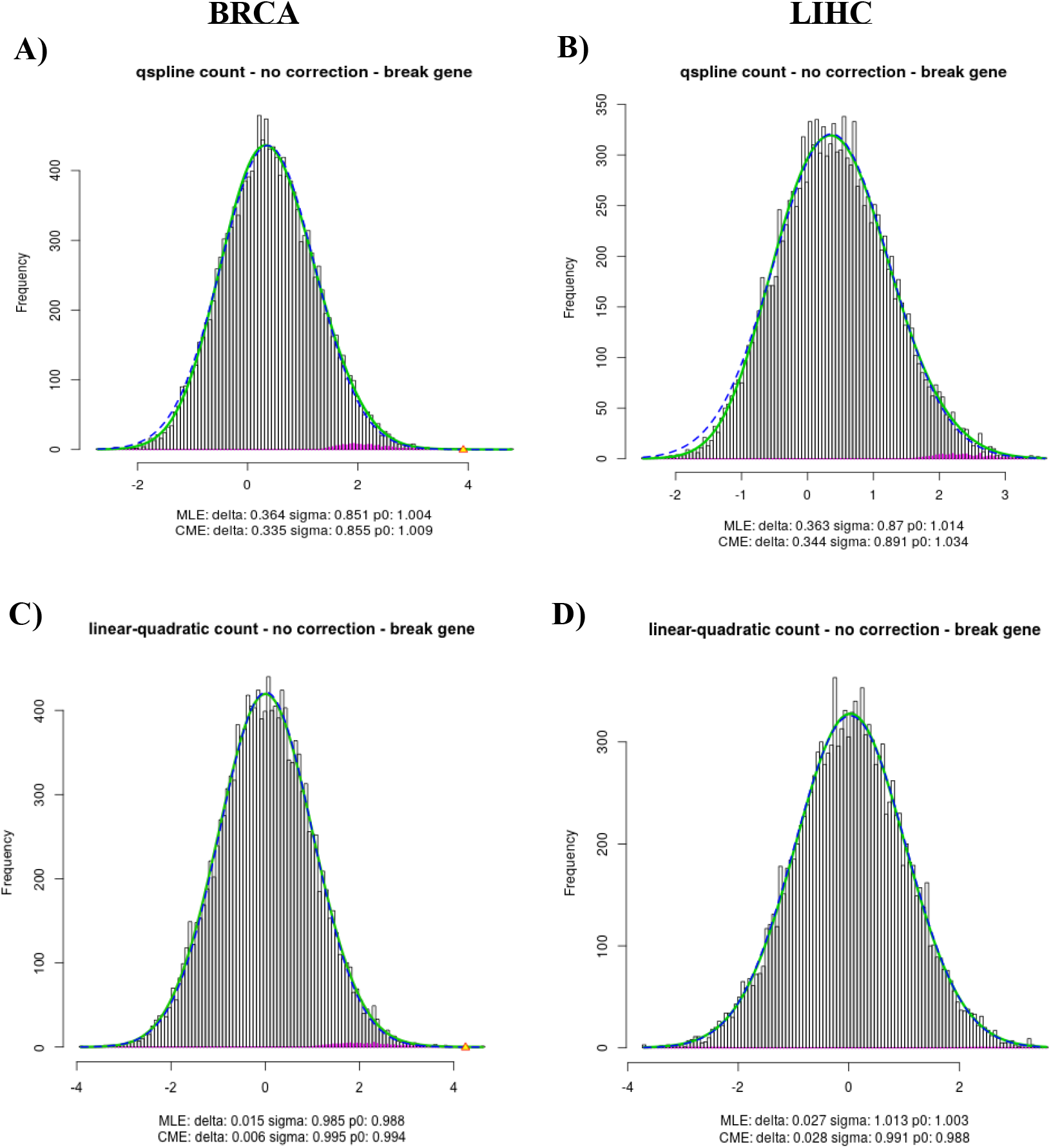
Z-scores for the association between gene expression and survival on the permuted BRCA (left panels) an LIHC (right panels) datasets. The blue dashed curves represent the estimated empirical null component densities. The green solid curves represent the fitted mixture density, i.e. a mixture between the null genes and marker genes (non-null). The data were permuted according to the first permutation strategy, as described in Figure S9. **Panels A and B; Z-score distributions when modelling gene expression with a spline function, ignoring confounders**. For both permuted datasets, the observed empirical null distribution is shifted and narrowed with respect to the theoretical null, suggesting failed mathematical assumptions. **Panels C and D; Z-score distributions when modelling gene expression with a linear and quadratic term, ignoring confounders**. With the linear and quadratic term for gene expression, we no longer observe deviations from the theoretical null, suggesting that the mathematical assumptions hold for this model.

As we observed failure of mathematical assumptions for the spline-based model but not for a model with linear and quadratic terms for gene expression, we continue exploring the other three issues for deviations of the theoretical null distribution for the latter model.

To assess the issue of correlation across genes, we generate data under a second permutation strategy (Figure S11). Here we will randomly permute the survival data of the different patients. This breaks the association (i) between a patient’s gene expression profile and survival, and (ii) between a patient’s confounders (i.e. both observed and unobserved confounders) and survival. Note, that the correlation across genes are retained in this scenario, as opposed to the previous permutation strategy.

**Figure S11:**
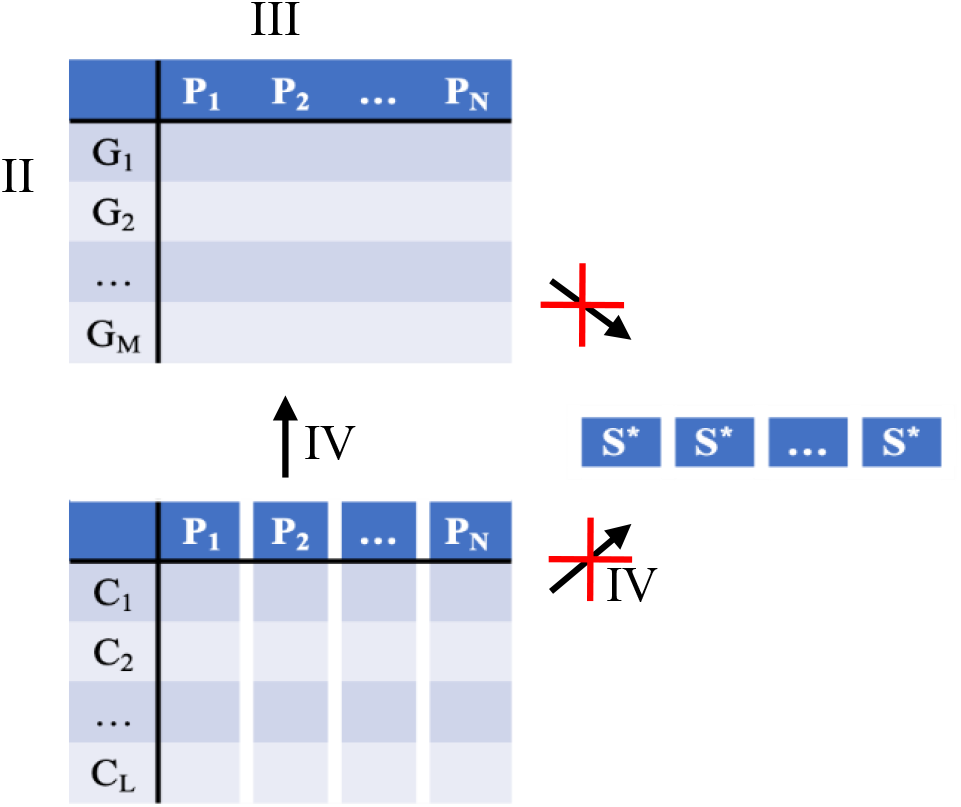
Second permutation strategy. We randomly permute the patient survival data, as indicated by the asterisk in the survival vector. We repeat the permutation six times for both the BRCA and LIHC datasets so that the conclusions are not driven by one specific permutation. The permutation breaks the association (i) between a patient’s gene expression profile and survival data, and (ii) between a patient’s confounders (i.e. both observed and unobserved confounders) and survival data. Comparing with the first permutation strategy shown in Figure S9, this second permutation strategy allows to assess the effect of issue II.

The empirical null distributions for a permuted dataset from each case study are depicted in Figure S12.

**Figure S12:**
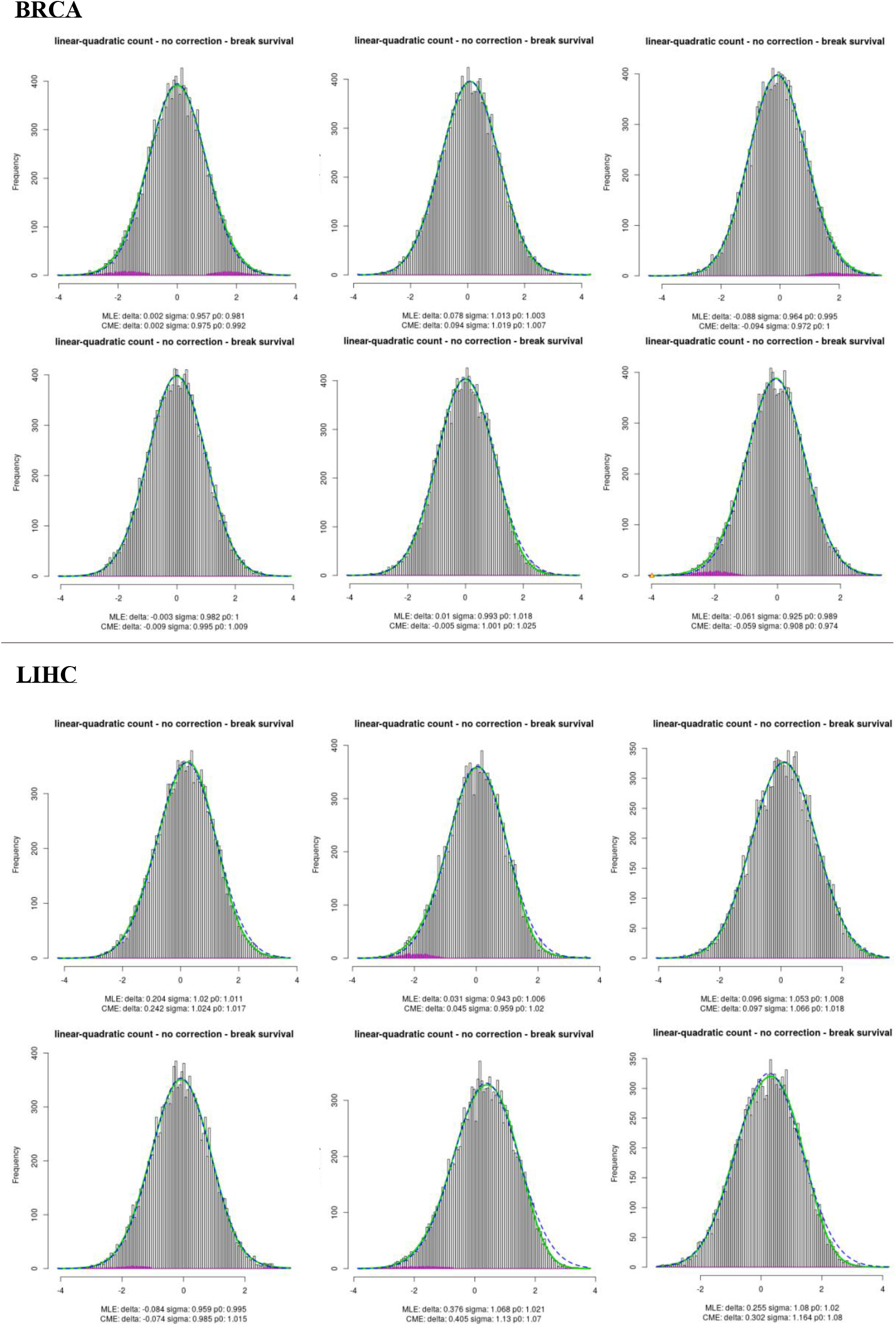
Z-scores for the association between gene expression and survival on six permuted BRCA (top six panels) an LIHC (bottom six panels) datasets. The blue dashed curves represent the estimated empirical null component densities. The green solid curves represent the fitted mixture density, i.e. a mixture between the null genes and marker genes (non-null). The data were permuted according to the second permutation strategy, as described in Figure S11. To make sure that are conclusions are not driven by one specific permutation of the survival data, we repeated the permutation six times for each dataset. For the permuted BRCA dataset, the estimated empirical null distribution overlaps almost perfectly with the theoretical standard normal distribution (Δ; [−0.09; 0.08] and σ; [0.93; 1.01] for six repeated permutations) with only very minor deviations between the empirical null and the observed z-scores. For the LIHC dataset, we observe that the empirical null is shifted as compared to the theoretical null (Δ; [−0.08; 0.38] and σ; [0.94; 1.08]) for the six repeated permutations, with some deviations between the empirical null and the observed z-scores in the right tail.

We observe a clear discrepancy between the empirical null distribution in the permuted BRCA dataset and the permuted LIHC dataset. For the former, the estimated empirical null distribution overlaps almost perfectly with the theoretical standard normal distribution (Δ; [−0.09; 0.08] and σ; [0.93; 1.01] for six repeated permutations) with only very minor deviations between the empirical null and the observed z-scores. This suggests that for the BRCA dataset that issue II, potential correlation across genes, does not cause major problems in the BRCA case study. For the LIHC dataset, we observe that the empirical null is shifted as compared to the theoretical null (Δ; [−0.08; 0.38] and σ; [0.94; 1.08]) for the six repeated permutations, with some deviations between the empirical null and the observed z-scores in the right tail. This suggests issue II is one of the underlying problems in the LIHC dataset.

In Figure S8, panels E and F, we showed that for both the BRCA and LIHC case studies the empirical null distributions are shifted and widened as compared to the theoretical standard normal distribution, when modeling survival as a function of gene expression with a linear and a quadratic term. This analysis, however, did not yet correct for the measured confounders. Therefore, we here inspect the empirical null distribution for both case studies when we additionally include a correction for the available baseline confounders through a spline function (age for the BRCA dataset, age and BMI for LIHC dataset). Note, that while we advised against modelling gene expression with a spline function of fixed degrees of freedom, modelling the confounders with a spline model should not be problematic, as we are not evaluating significance of the spline term in the actual inference (with the likelihood ratio test we only assess gene expression and leave the spline for age in the model). The empirical null distributions of this analysis are depicted in Figure S13.

For both analyses we observe that the empirical null distribution remains shifted (Δ = 0.448 for BRCA and Δ = 1.036 for LIHC) and wider (σ = 1.157 for BRCA and σ = 1.465 for LIHC) than the theoretical null, similar to the results obtained in Figure S8, panels E and F. This is in an indication that reason (IV) unmeasured confounders is one of the main drivers in the failure of the theoretic null.

Initially, we proposed semi-parametric CPH model that studies the association between gene expression and survival while correcting for baseline confounders (age for the BRCA dataset, age and BMI for the LIHC dataset) with a spline function. As shown above, this model suffers from all four issues discussed by Efron (*5*) regarding the theoretical null distribution of the test statistics, jeopardizing correct statistical inference. However, when replacing the spline-based LRT test with an LRT test based on a linear and quadratic GE-effect, we no longer observed failure of the null due to mathematical assumptions (Figure S10). Nevertheless, the null of this quadratic CPH model still suffers from unobserved confounders (BRCA and LIHC, Figure S13) and from correlation between genes (LIHC, Figure S12). The former, however, could not be addressed using the publicly available baseline confounders.

**Figure S13:**
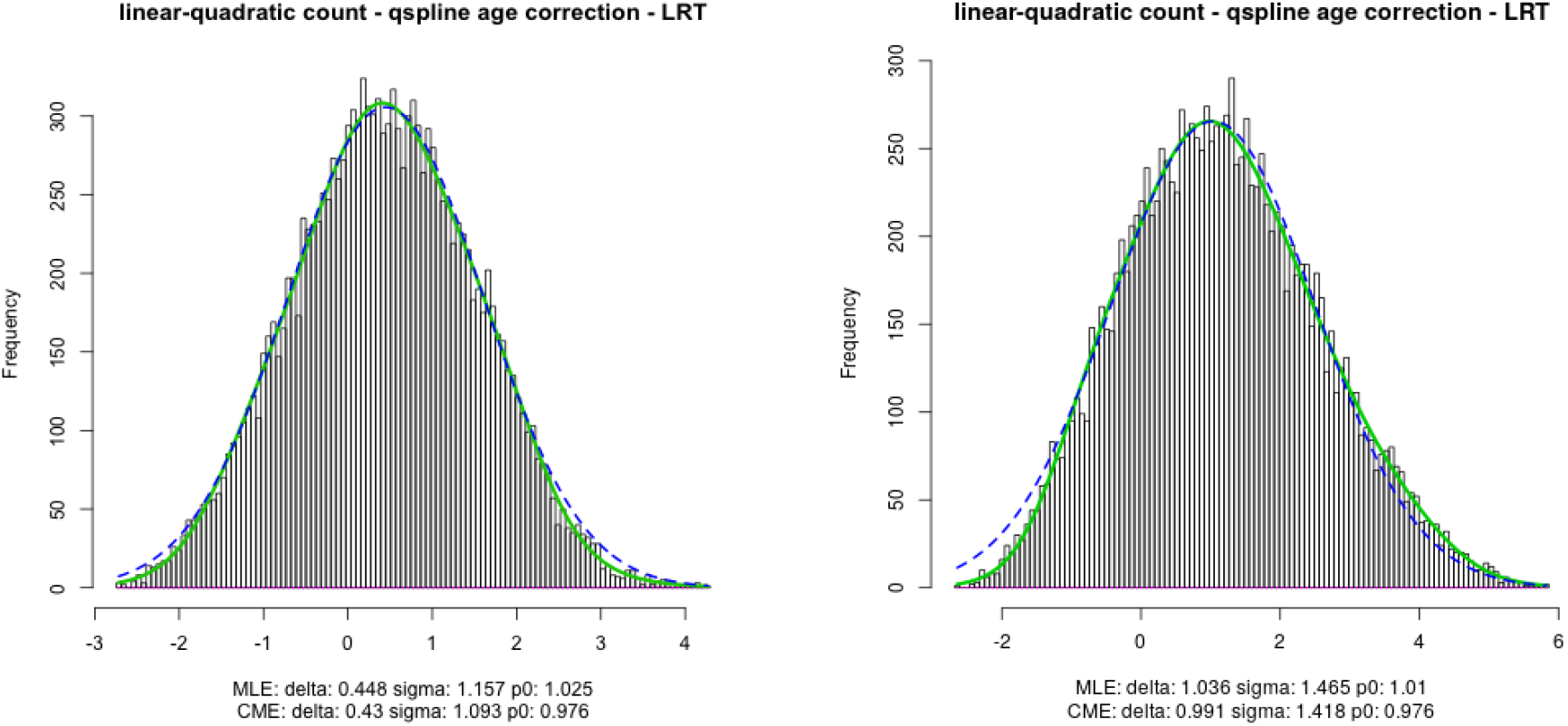
Z-scores for the association between gene expression and survival on the original BRCA (left panel) an LIHC (right panel) datasets when modelling gene expression with a linear and quadratic term and correcting for confounders with a spline function. The blue dashed curves represent the estimated empirical null component densities. The green solid curves represent the fitted mixture density, i.e. a mixture between the null genes and marker genes (non-null). For both analyses we observe that the empirical null distribution is shifted (Δ = 0.448 for BRCA and Δ = 1.036 for LIHC) and wider (σ = 1.157 for BRCA and σ = 1.465 for LIHC) than the theoretical null. This is in an indication that reason (IV) unmeasured confounders is one of the main drivers in the failure of the theoretic null. Note that the empirical null distribution does not very well fit the observed z-scores.

While the empirical null distribution of our final inference procedure did not perfectly fit the observed z-scores (Figure S13), we feel that this is the current state-of-the-art workflow to analyze the publicly available datasets.

## 5. Top list of genes associated with survival

There are two major conclusions to our analysis with respect to the discovery of marker genes. First, correcting for confounders has an impact on the ranking of the genes with respect to their association with survival. This indicates that correcting for confounders in this type of analysis is important. This is further emphasized by the fact that, as shown in table S2, two known genes in breast cancer (MMP13 and RGL3) are in the top 10 most significantly associated genes only in our improved workflow. Second, when the p-values obtained through our final CPH model were corrected for (i) deviations from the underlying theoretical null distribution and (ii) multiple testing through the lfdr procedure proposed by Efron (*6*, *7*), there was no longer any statistical evidence for marker genes that are associated with survival in the BRCA and LIHC case studies (Note, that according to the guidelines of Efron, a significance level of 0.2 is used for the lfdr). This suggests that the original marker genes that were proposed by Uhlen et al. (*1*) and that are still being communicated via the Human Pathology Atlas web resource can be expected to contain a vast number of false positive markers. In tables S1–4, we display the top 10 marker genes that are associated with survival in the BRCA and LIHC datasets according to the original (Tables 1 and 3) and our updated workflow (Tables 2 and 4). The complete tables may be retrieved from our companion GitHub page (/dataBRCA and /dataLIHC).

---

**Table S1:** **Effect of our proposed step-by-step improvement of the data analysis workflow on the significance levels of the top 10 prognostic genes in breast cancer as communicated by Uhlen et al (1).** In total, 14.573 genes were assessed. Note, that *CYP24A1* was not included in our analysis, as the expression level of this gene was below the adopted threshold. **Column 1**: the p-value for each gene, as obtained with the CPH model, is displayed. **Column 2**: the p-values are obtained again by CPH models, that additionally account for the confounder age. **Column 3**: the p-values of the second column were corrected for multiple testing with the FDR method of Benjamin and Hochberg *(3)*. **Column 4**: the p-values of the second column were corrected for (i) deviations from the underlying theoretical null distribution and (ii) multiple testing through Efron’s lfdr procedure *(6,7)*. **Column 5**: the p-values as communicated in the original publication and the Human Pathology Atlas web resource. **Column 6**: Prognostic type of the gene as determined by the original publication. Can be either favorable or unfavorable. **Column 7**: the rank of the initial top 10 genes based on our workflow (based on the CPH age corrected p-values). Note, that the ordering has changed dramatically for some of the genes, which is mainly due to our correction for the confounder age. **Column 8**: the rank of the initial top 10 genes based on the original analysis of Uhlen et al. 2017 using the Kaplan-Meier p-values.

| Symbols | Gene only | Gene corrected | Gene corrected BH-FDR | Gene Corrected lfr | Kaplan Meier | Prognostic Types | Rank our | Rank Uhlen |
| --- | --- | --- | --- | --- | --- | --- | --- | --- |
| LRP11 | 2.49E-04 | 5.67E-04 | 1.40E-01 | 1 | 2.31E-08 | Unfav. | 52 | 1 |
| ZNF385B | 6.23E-05 | 1.07E-04 | 1.40E-01 | 1 | 4.86E-08 | Fav. | 9 | 2 |
| PGK1 | 7.49E-06 | 3.70E-05 | 1.35E-01 | 1 | 7.05E-08 | Unfav. | 4 | 3 |
| CYP24A1 | NA | NA | NA | NA | 1.19E-07 | Fav. | NA | 4 |
| PCMT1 | 6.19E-05 | 6.04E-04 | 1.40E-01 | 1 | 4.02E-07 | Unfav. | 56 | 5 |
| TP53AIP1 | 6.14E-06 | 6.25E-05 | 1.40E-01 | 1 | 4.12E-07 | Fav. | 5 | 6 |
| TFPI2 | 1.90E-04 | 4.90E-03 | 2.08E-01 | 1 | 8.33E-07 | Fav. | 341 | 7 |
| FAM173B | 6.45E-03 | 2.35E-02 | 2.76E-01 | 1 | 1.32E-06 | Unfav. | 1237 | 8 |
| IL27RA | 3.57E-04 | 5.25E-03 | 2.10E-01 | 1 | 1.74E-06 | Fav. | 362 | 9 |
| JCHAIN | 6.22E-04 | 4.07E-03 | 1.96E-01 | 1 | 1.81E-06 | Fav. | 299 | 10 |

**Table S2:** Effect of our proposed step-by-step improvement of the data analysis workflow on the significance levels of the top 10 prognostic genes in the breast cancer dataset as obtained with our workflow. In total, 14.573 genes were assessed. Note, that two known genes in breast cancer (MMP13 and RGL3) are in the top 10 most significantly associated genes only in our improved workflow. **Column 1**: the p-value for each gene, as obtained with the CPH model, is displayed. **Column 2**: the p-values are obtained again by CPH models, that additionally account for the confounder age. **Column 3**: the p-values of the second column were corrected for multiple testing with the FDR method of Benjamin and Hochberg *(3)*. **Column 4**: the p-values of the second column were corrected for (i) deviations from the underlying theoretical null distribution and (ii) multiple testing through Efron’s lfdr procedure *(6,7)*. **Column 5**: the p-values as communicated in the original publication and the Human Pathology Atlas web resource. **Column 6**: Prognostic type of the gene as determined by the original publication. Can be either favorable or unfavorable. **Column 7**: the rank of the initial top 10 genes based on our workflow (based on the CPH age corrected p-values). Note, that the ordering has changed dramatically for some of the genes, which is mainly due to our correction for the confounder age. **Column 8**: the rank of the initial top 10 genes based on the original analysis of Uhlen et al. 2017 using the Kaplan-Meier p-values.

| Symbols | Gene only | Gene corrected | Gene corrected BH-FDR | Gene corrected lfr | Kaplan Meier | Prognostic Types | Rank our | Rank Uhlen |
| --- | --- | --- | --- | --- | --- | --- | --- | --- |
| RGL3 | 4.94E-05 | 2.55E-05 | 1.35E-01 | 1 | 7.35E-04 | Fav. | 1 | 698 |
| SERPINA1 | 1.94E-04 | 2.67E-05 | 1.35E-01 | 1 | 1.61E-05 | Fav. | 2 | 52 |
| TTC39C | 6.20E-04 | 3.07E-05 | 1.35E-01 | 1 | 1.48E-04 | Fav. | 3 | 264 |
| PGK1 | 7.49E-06 | 3.70E-05 | 1.35E-01 | 1 | 7.05E-08 | Unfav. | 4 | 3 |
| TP53AIP1 | 6.14E-06 | 6.25E-05 | 1.40E-01 | 1 | 4.12E-07 | Fav. | 5 | 6 |
| MMP13 | 1.26E-03 | 6.30E-05 | 1.40E-01 | 1 | 8.98E-03 | Unfav. | 6 | 3042 |
| QPRT | 2.27E-04 | 6.90E-05 | 1.40E-01 | 1 | 5.10E-05 | Unfav. | 7 | 127 |
| TRAPPC12 | 1.30E-03 | 1.06E-04 | 1.40E-01 | 1 | 4.86E-08 | Fav. | 8 | 2 |
| ZNF385B | 6.23E-05 | 1.07E-04 | 1.40E-01 | 1 | 1.80E-05 | Fav. | 9 | 57 |
| PARP3 | 5.62E-03 | 1.09E-04 | 1.40E-01 | 1 | 6.39E-05 | Fav. | 10 | 144 |

**Table S3:** **Effect of our proposed step-by-step improvement of the data analysis workflow on the significance levels of the top 10 prognostic genes in liver cancer as communicated by Uhlen et al (1).** In total, 13.376 genes were assessed. **Column 1**: the p-value for each gene, as obtained with the CPH model, is displayed. **Column 2**: the p-values are obtained again by CPH models, that additionally account for confounders age and BMI. **Column 3**: the p-values of the second column were corrected for multiple testing with the FDR method of Benjamin and Hochberg *(3)*. **Column 4**: the p-values of the second column were corrected for (i) deviations from the underlying theoretical null distribution and (ii) multiple testing through Efron’s lfdr procedure *(6,7)*. **Column 5**: the p-values as communicated in the original publication and the Human Pathology Atlas web resource. **Column 6**: Prognostic type of the gene as determined by the original publication. Can be either favorable or unfavorable. **Column 7**: the rank of the initial top 10 genes based on our workflow (based on the CPH age and BMI corrected p-values). **Column 8**: the rank of the initial top 10 genes based on the original analysis of Uhlen et al. 2017 using the Kaplan-Meier p-values.

| Symbols | Gene only | Gene corrected | Gene Corrected BH-FDR | Gene Corrected lfr | Kaplan Meier | Prognostic Types | Rank our | Rank Uhlen |
| --- | --- | --- | --- | --- | --- | --- | --- | --- |
| SFPQ | 1.06E-06 | 1.25E-07 | 8.63E-05 | 1 | 7.91E-13 | Unfav. | 18 | 1 |
| G6PD | 2.83E-07 | 3.21E-08 | 4.10E-05 | 1 | 9.45E-13 | Unfav. | 10 | 2 |
| KIF20A | 8.02E-07 | 7.17E-08 | 6.39E-05 | 1 | 2.14E-12 | Unfav. | 15 | 3 |
| KDM1A | 3.68E-07 | 9.66E-09 | 3.22E-05 | 1 | 4.47E-12 | Unfav. | 3 | 4 |
| TRMT6 | 2.09E-08 | 7.15E-09 | 3.22E-05 | 1 | 4.70E-12 | Unfav. | 1 | 5 |
| PSMD1 | 1.61E-06 | 1.61E-07 | 9.40E-05 | 1 | 8.63E-12 | Unfav. | 22 | 6 |
| GTPBP4 | 1.28E-06 | 2.07E-06 | 3.38E-04 | 1 | 8.85E-12 | Unfav. | 81 | 7 |
| CCT4 | 2.14E-08 | 1.75E-08 | 3.34E-05 | 1 | 9.37E-12 | Unfav. | 7 | 8 |
| CAD | 2.27E-06 | 3.20E-07 | 1.22E-04 | 1 | 1.72E-11 | Unfav. | 35 | 9 |
| CBX2 | 6.13E-08 | 8.77E-08 | 7.34E-05 | 1 | 2.25E-11 | Unfav. | 16 | 10 |

**Table S4:** Effect of our proposed step-by-step improvement of the data analysis workflow on the significance levels of the top 10 prognostic genes in the liver cancer dataset as obtained with our workflow. In total, 13.376 genes were assessed. **Column 1**: the p-value for each gene, as obtained with the CPH model, is displayed. **Column 2**: the p-values are obtained again by CPH models, that additionally account for confounders age and BMI. **Column 3**: the p-values of the second column were corrected for multiple testing with the FDR method of Benjamin and Hochberg *(3)*. **Column 4**: the p-values of the second column were corrected for (i) deviations from the underlying theoretical null distribution and (ii) multiple testing through Efron’s lfdr procedure *(6,7)*. **Column 5**: the p-values as communicated in the original publication and the Human Pathology Atlas web resource. **Column 6**: Prognostic type of the gene as determined by the original publication. Can be either favorable or unfavorable. **Column 7**: the rank of the initial top 10 genes based on our workflow (based on the CPH age and BMI corrected p-values). **Column 8**: the rank of the initial top 10 genes based on the original analysis of Uhlen et al. 2017 based on the Kaplan-Meier p-values.

| Symbols | Gene only | Gene corrected | Gene corrected BH-FDR | Gene corrected lfr | Kaplan Meier | Prognostic Types | Rank our | Rank Uhlen |
| --- | --- | --- | --- | --- | --- | --- | --- | --- |
| TRMT6 | 2.09E-08 | 7.15E-09 | 3.22E-05 | 1 | 2,30E-10 | Unfav. | 1 | 5 |
| PSRC1 | 1.86E-07 | 9.49E-09 | 3.22E-05 | 1 | 2,14E-12 | Unfav. | 2 | 31 |
| KDM1A | 3.68E-07 | 9.66E-09 | 3.22E-05 | 1 | 4,70E-12 | Unfav. | 3 | 4 |
| KPNA2 | 7.88E-07 | 9.86E-09 | 3.22E-05 | 1 | 2,12E-10 | Unfav. | 4 | 18 |
| HDAC2 | 9.02E-08 | 1.21E-08 | 3.22E-05 | 1 | 7,44E-08 | Unfav. | 5 | 22 |
| CCDC77 | 1.44E-06 | 1.72E-08 | 3.34E-05 | 1 | 5,39E-10 | Unfav. | 6 | 373 |
| CCT4 | 2.14E-08 | 1.75E-08 | 3.34E-05 | 1 | 6,45E-10 | Unfav. | 7 | 8 |
| UTP11L | 3.66E-08 | 2.12E-08 | 3.54E-05 | 1 | 4,30E-11 | Unfav. | 8 | 163 |
| HILPDA | 3.98E-07 | 2.74E-08 | 4.07E-05 | 1 | 1,36E-09 | Unfav. | 9 | 12 |
| G6PD | 2.83E-07 | 3.21E-08 | 4.10E-05 | 1 | 9,19E-09 | Unfav. | 10 | 2 |

## Notes

https://github.com/statOmics/pitfallsOfHumanPathologyAtlas

### Competing Interest Statement

The authors have declared no competing interest.

### Summary of Updates

Elaborate on results of mock analysis in the multiple testing paragraph, clarify captions of Figures S8-S13, minor adjustments in formulations across the main and supplementary texts.

